# Energy Landscapes from Cryo-EM Snapshots: A Benchmarking Study

**DOI:** 10.1101/2022.06.13.495454

**Authors:** Raison Dsouza, Ghoncheh Mashayekhi, Roshanak Etmaadpour, Peter Schwander, Abbas Ourmazd

## Abstract

Biomolecules undergo complex continuous conformational motions, a subset of which are functionally relevant. Understanding, and ultimately controlling biomolecular function are predicated on the ability to map continuous conformational motions and identify the functionally relevant conformational trajectories. For equilibrium and near-equilibrium processes, the function proceeds along minimum-energy pathways on one or more energy landscapes, because higher-energy conformations are only weakly occupied. With the growing interest in identifying functional trajectories on energy landscapes, the reliable mapping of energy landscapes has become paramount. In response, various data-analytical tools for determining structural variability are emerging. A key question concerns the veracity with which each data-analytical tool can extract functionally relevant conformational trajectories from a collection of singleparticle cryo-EM snapshots. Using synthetic data as an independently known ground truth, we benchmark the ability of four leading algorithms to determine biomolecular energy landscapes and identify the functionally relevant conformational paths on these landscapes. Such benchmarking is essential for systematic progress toward atomic-level movies of continuous biomolecular function.

---

Biomolecular machines have evolved to perform specific tasks through a concerted sequence of conformational motions. There is growing recognition that such motions involve continuous conformational changes, rather than jumps between a small number of discrete states [1, 2]. Apart from disordered proteins, conformational continua span a spectrum of different energies. In thermal equilibrium, the probability of a conformational state being occupied is determined by the Boltzmann factor, which drops exponentially with increasing energy.

Conformational motions of proteins can thus be represented as low-lying (and thus strongly occupied) pathways on one or more energy landscapes (EL) [3, 4]. In principle, an unlimited number of conformational paths connect a “start” conformation A to an “end” conformation B. However, most such paths include high-energy states, which are sparsely populated under biologically relevant conditions. Due to the exponential nature of the Boltzmann inverse relationship between energy and occupation probability [5], lowest-energy conformational paths contribute maximally to function.

The growing recognition of the importance of energy landscapes for discerning function has spawned an increasing number of sophisticated algorithms capable of mapping continuous conformational motions. Using a synthetic dataset of cryo-EM snapshots with known ground truth energy landscape, we compare the performance of four leading algorithms, specifically Relion Multibody [6], CryoSPARC 3DVA [7], Manifold-EM [8], and CryoDRGN VAE [9] in faithfully extracting the energy landscape from snapshots. We benchmark the performance of each algorithmic approach in terms of the accuracy with which the underlying energy landscape is recovered from the data.

To date, no comparative benchmarking study of the strengths and weaknesses of different data-analytical approaches for analyzing continuous conformations has been reported. The lack of comparative benchmarks seriously hampers the assessment of the usefulness and reliability of different algorithmic tools in extracting information from experimental data.

In this paper, we use a synthetic dataset generated with an a priori known “ground-truth” energy landscape to benchmark, in silico, the above four leading algorithms for conformational analysis of cryo-EM data. Specifically, we test each method’s ability to: (a) Recover the correct energy landscape from synthetic cryo-EM datasets; (b) Reveal the functionally important conformational degrees of freedom; and (c) Identify the functionally relevant conformational paths on these landscapes. Although the nature and number of potentially useful algorithms are currently evolving, the four selected approaches represent the state of the art in mapping continuous conformational motions from ensembles of single-particle cryo-EM snapshots.

The primary goals of this paper are therefore twofold:

i. Benchmark the performance of the four leading algorithms listed above in faithfully extracting conformational energy landscapes from synthetic cryo-EM snapshots; and,
ii. Provide a well-characterized synthetic cryo-EM dataset suitable for benchmarking, in order to facilitate the development of more effective data-analytical tools capable of identifying functionally relevant conformational landscapes and motions.

The synthetic dataset of cryo-EM snapshots (with noise SNR=1) pertains to a ribosome-like object with two conformational degrees of freedom, reflecting an underlying energy landscape of twelve energy minima (Figure 1(a)). The twelve energy minima are arranged in a 3×4 grid. The distribution of points reflects the underlying occupancy distribution, which is designed to have modulations to clearly indicate minima (stable states) and maxima (transition states) (Figure 1(b)). Each energy minimum has been assigned a unique color to identify each energy minimum. (Figure1(c)). The distribution of the ensemble of 3 million snapshots reflects the rotations of the small subunit (SSU) about two axes corresponding to conformational coordinates 1 and 2. The SSU of the ribosome-like object is permitted to rotate in a ratchet-like manner about two mutually orthogonal axes, with the large subunit (LSU) fixed (Figure 1(d)).

**Figure 1:**
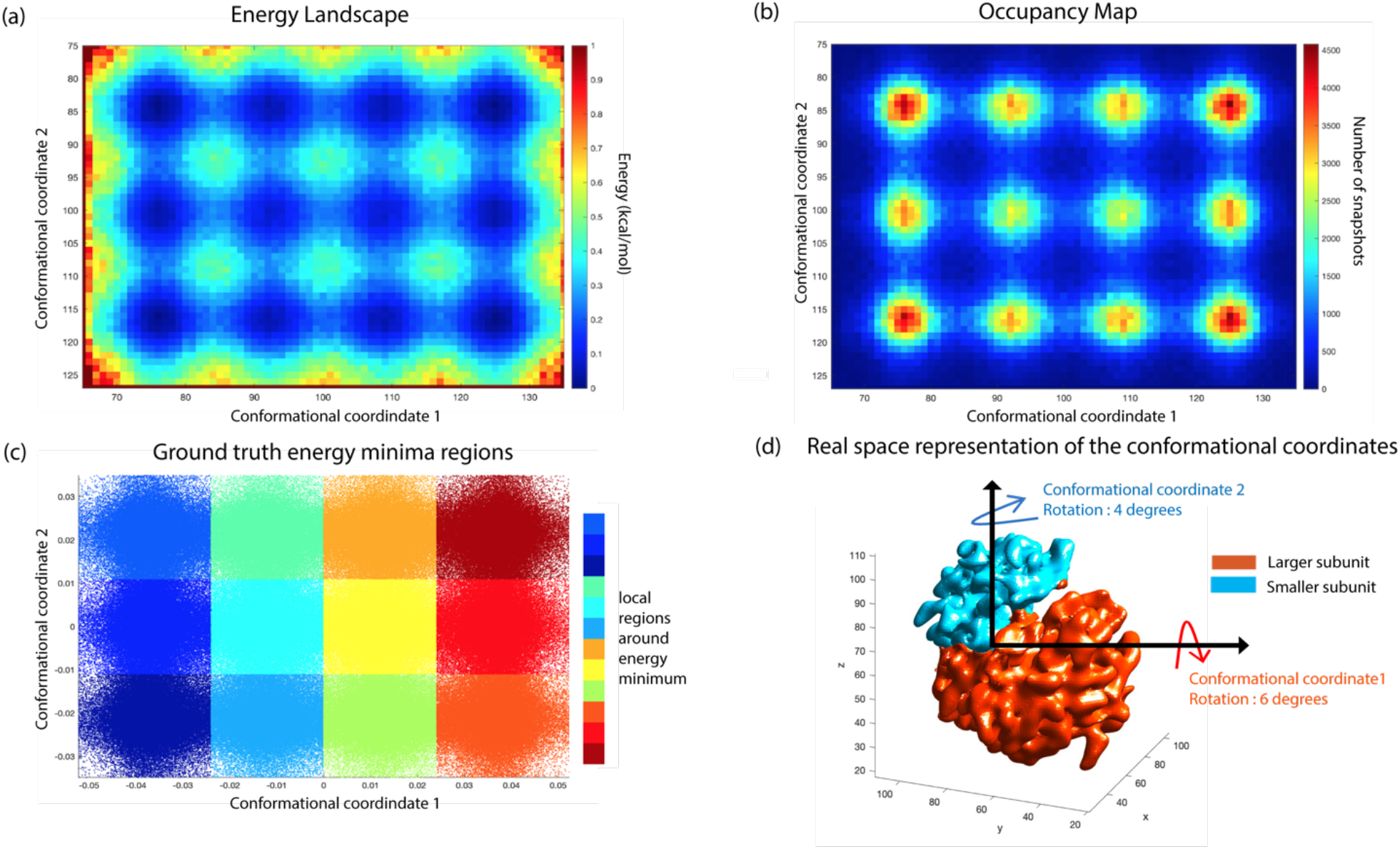
(a). The energy landscape is spanned by the two conformational coordinates with color bar representing the local density of snapshots. (b). The occupancy map of the conformational change along with the coordinates as a function of the number of snapshots (c). The ground truth energy landscape where the region around each minimum is colored differently (d).Sketch of the synthetic model, rotation of the small subunit about the two axes labeled conformational coordinates 1 and 2 respectively. The larger subunit (orange) is kept fixed.

The performance of each of the four data-analytical approaches is quantified in terms of the fidelity of the energy landscape recovered from the synthetic snapshots. The fidelity is quantified in terms of accuracy, defined as the distance between a given set of data points from their true value. Details are provided in the Methods section (Under Accuracy).

## RESULTS

Each of the four data-analytical software tools evaluated in this benchmarking study was applied to the above dataset. The ground-truth snapshot orientations were provided to each algorithm to focus the study on the ability of the four different algorithms to extract conformational information. From each algorithm, we analyzed the top two (i.e., the most “powerful”) conformational coordinates. We map the problem of analyzing the topology of the output energy landscape as the ability to classify snapshots correctly into the 12 ground truth energy minima. This allows us to use well-known metrics from classification.

### Accuracy of the energy landscapes extracted by each of the four data-analytical tools

We use Recall [10] defined as

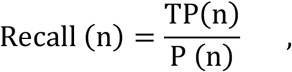

where P(n) is the so-called positive class (snapshots belonging to ground truth energy minima ‘n’) and TP(n) are the true positives i.e., snapshots correctly assigned by the algorithm to the energy minimum ‘n’. This allows us to compute a recall value for each of the twelve energy minima (See SI Tables 1-4).

The Average of Recall values also called Balanced Accuracy [11], or simply Accuracy allows us to assign a quantitative score to each algorithm’s ability to accurately extract the energy landscape underlying the synthetic data (See Methods Under Accuracy). Essentially one tracks the region of snapshots around the energy minimum in the input landscape and asks, “What is the extent of deformation of the energy minima after the application of the algorithm?”. To answer this, we have used the Euclidean metric (L2-norm) to calculate the neighborhood of points in both the input and output (Figure 2(a)). The accuracy of each method is given in Table 1.

**Figure 2:**
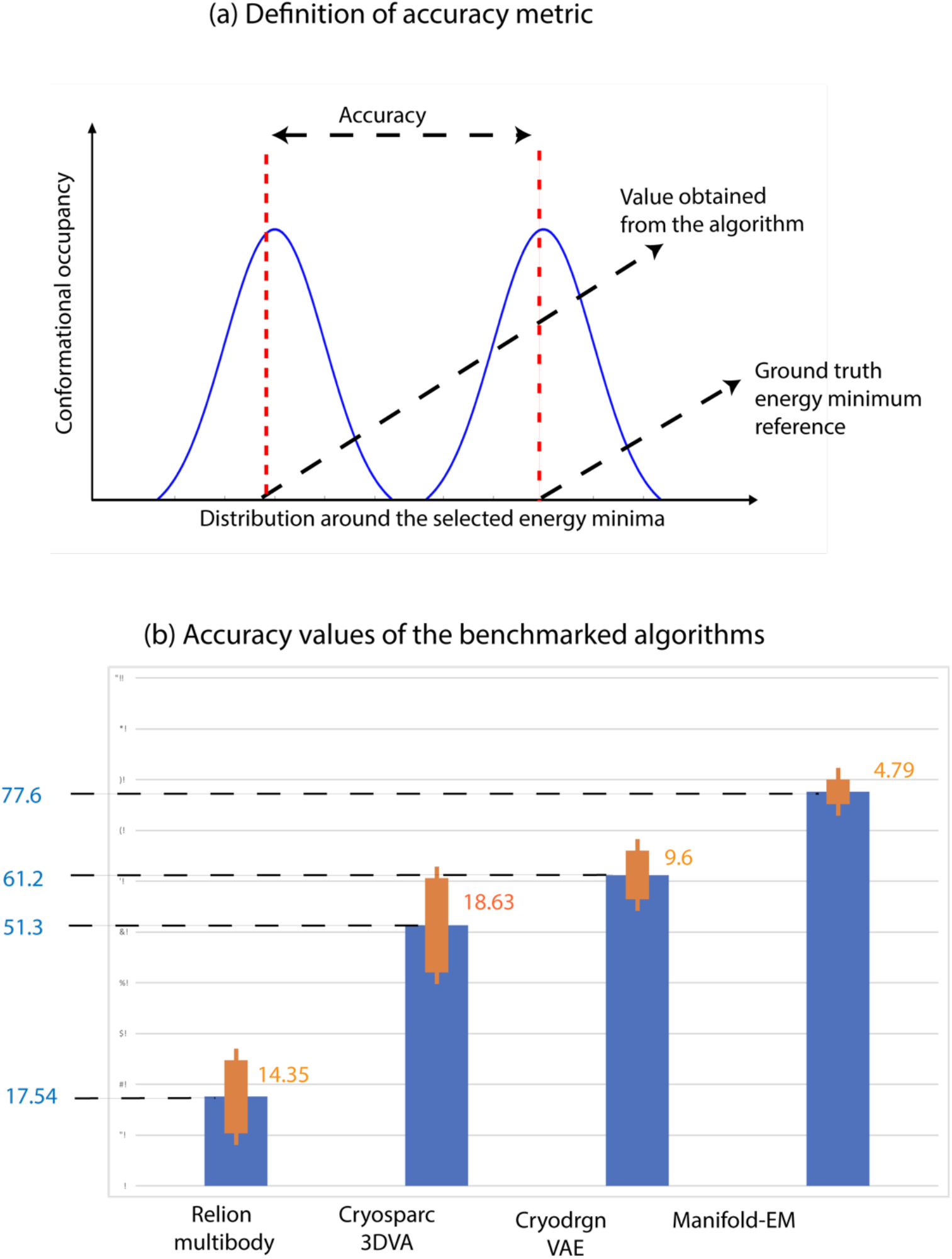
(a). A pictorial description of the recall metric calculation in context of energy landscape benchmarking. (b). Bar plots containing accuracy values obtained from each method where ground truth information was provided.

**Table 1.**
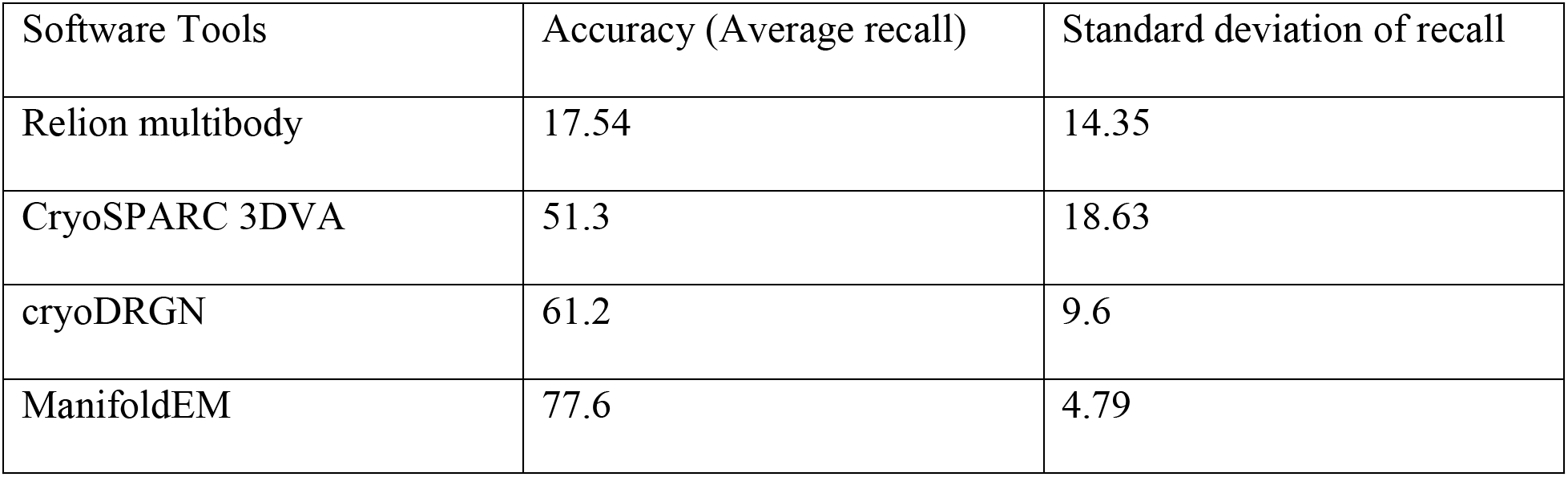
Statistics of the Accuracy metric for the various algorithms used in the benchmarking

| Software Tools | Accuracy (Average recall) | Standard deviation of recall |
| --- | --- | --- |
| Relion multibody | 17.54 | 14.35 |
| CryoSPARC 3DVA | 51.3 | 18.63 |
| cryoDRGN | 61.2 | 9.6 |
| ManifoldEM | 77.6 | 4.79 |

In brief, the Relion multibody assigns only about 17.54 ± 14.35% of the points on average to the correct region in the energy landscape (Figure 2(b)). (Accuracy is averaged over 12 energy minima). The cryoSPARC 3DVA algorithm shows an accuracy score of 51.3 ± 18.63%.cryoDRGN variational autoencoder assigns snapshots with an accuracy of 61.2 ± 9.6%. The Manifold-EM algorithm correctly assigns 77.6 ± 4.79 % of the snapshots on average over the entire ground truth energy landscape. Among the investigated approaches, the accuracy of Manifold-EM represents the highest score and the smallest uncertainty.

### Occupancy maps and energy landscapes

The occupation probability of a conformational state is determined by the energy of the state via the Boltzmann factor. An occupancy map is simply an alternative representation of the conformational energy landscape of the ribosome-like object. Figures 1(c) and 1(d) show the occupancy map and energy landscapes for the conformational states of the synthetic ribosome.

To date, Manifold EM is the only method that can calculate the occupancy map of conformations and energy landscapes. The occupancy map and the corresponding energy landscape obtained from the Manifold-EM data analytical pipeline are shown in Figures 2(a) and 2(b) respectively. It is evident from the occupancy map (Figure 2(a)) that Manifold-EM not only captures the high occupancy regions or equivalently twelve energy minimum regions (Figure 2(b)), but also preserves the modulated features of the occupancy across the ground truth conformational coordinates (Figure 1(c)). The other benchmarked algorithms have not yet developed a way to extract thermodynamic quantities like occupancy probability or free energy.

### Visualizing the energy landscape

To further support the claims from the accuracy metric, we looked at the distribution of data points in the output using the ground truth energy minima as 12 categories or classes. This procedure is known as ‘parentage’ or ‘lineage’ [12]. Essentially, parentage uses the ground truth labels for each single-particle snapshot in the energy landscape and identifies its position in the output of each of the data-analytical tools (Figures 3(a-d)). The region surrounding each of the 12-energy minima is rendered in a different color to elucidate the extent to which snapshots stemming from adjacent regions of the ground truth energy landscape are correctly assigned by each of the four data-analytical techniques.

**Figure 3:**
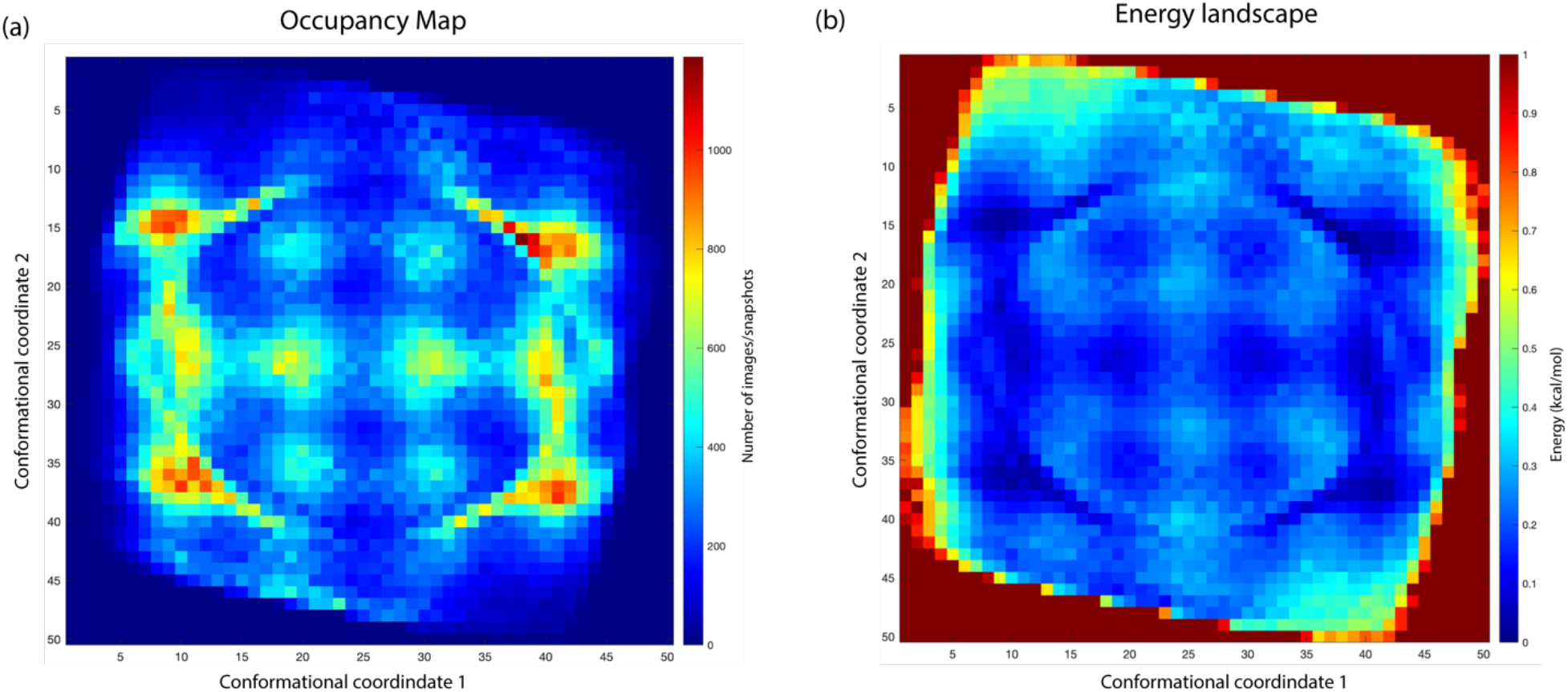
(a). Occupancy map and (b). Energy landscape obtained by the data-analytical pipeline implemented in Manifold-EM.

As shown in Figure 3(a-d), the overall ground-truth topology is preserved to the largest extent by ManifoldEM. Relion, cryoSPARC, and cryoDRGN severely distort the energy landscape, intermixing points stemming from different energy minima. When the snapshots are represented using a two-dimensional representation (in terms of the two conformational coordinates), the different algorithms show distinctive distributions of the input snapshots. In the lower-dimensional representation (2D), the points are now well separated in the case of Relion. They lie on top of each other. In cryoSPARC and cryoDRGN, about four such regions can be seen at once. The others are superimposed in the 2D space and not resolved. However, only Manifold EM correctly reduces the high-dimensional problem (image space) to the true low dimensional distribution (2D) of the ground truth.

## DISCUSSIONS

Inferring biological function from the structure is a paramount goal of structural biology. The results presented here highlight the importance of basing functional inference on energy landscapes and conformational coordinates derived from the data. To this end, we have developed and tested a rigorous approach to benchmark the performance of four data-analytical algorithms in faithfully extracting energy landscapes underlying a collection of synthetic snapshots. This assessment is based on the accuracy with which individual snapshots are placed on the output energy landscape as compared with the independently known ground-truth distribution. Using a synthetic model consisting of a 3×4 array of 12 energy minima, Manifold-EM, recovers the underlying energy landscape most accurately. Our approach thus offers a rigorous means for measuring and improving the performance of data-analytical approaches capable of extracting energy landscapes from cryo-EM datasets.

The study design, including the synthetic ribosome-like object, was chosen to mimic conditions typically encountered in conformational motions in biological macromolecules. Of course, any benchmarking exercise pertains to a particular time point. Consistent with the rapid progress in cryo-EM single-particle imaging, data-analytical software tools are evolving apace [13–17]. Our approach offers a quantitative method for assessing and guiding this rapidly progressing field.

The recall and accuracy metric are powerful tools for comparing the different algorithms, as it allows the study of energy regions without prior assumptions about the shape and distribution of the energy landscapes. Despite differences in the formulation of the different algorithms, the accuracy-based score can be consistently applied.

According to our study, Manifold EM is currently the most robust way to extract energy landscapes and conformational coordinates from single-particle cryo-EM images. The method preserves about 77% of the snapshots in each energy minima and shows the least deviation of accuracy scores across the 12 minima (4.78%).

For future work, it will be powerful to complement with reconstructed molecular movies from the energy landscape with atomic coordinates at different resolutions as obtained from each method. We anticipate that the synthetic data which incorporates continuous conformations will become an important tool for algorithm validation.

## Author Contributions

RD wrote the paper, co-designed the benchmark study, and performed the calculations using the Synthetic Data. GM and RE performed the calculations for the Synthetic data using Manifold-EM. PS co-designed the study, helped analyze the results and generated the synthetic data for the study. AO co-designed the study, helped analyze the results in terms of the concepts of singleparticle cryo-EM and the dynamics of molecular machines, and co-wrote the paper.

RD, GM, PS, and AO analyzed and interpreted the results.

## Funding Sources

The development of underlying techniques was supported by the US Department of Energy, Office of Science, Basic Energy Sciences under award DE-SC0002164 (underlying dynamical techniques), and by the US National Science Foundation under awards STC-1231306 (underlying data analytical techniques) and DBI-2029533 (underlying analytical models).

## Acknowledgments

We acknowledge valuable discussions with current members of the Ourmazd group: Dr. Ahmad Hosseinizadeh, Dr. Russell Fung, and Dr. Eduardo Chu-Cruz.The research was conducted at the University of Wisconsin-Milwaukee.

## METHODS

### Accuracy metric

Finding an appropriate metric for the synthetic ribosome dataset is a challenge since the energy landscape is continuous and it is hard to quantify the distortion caused by the individual algorithms. We tackle this problem as a multi-class classifier, as the goal is to bin each particle in one of the 12 energy minima. The recall metric was implemented as follows:

1. The center of each energy minima was obtained by calculating the ‘mean’ of all the particles corresponding to that minima region (for all 12 regions)
2. Compute the distance matrix (squared Euclidean) using the 12 centers
3. Using the distance of each particle from the corresponding center (see Figure 4(a)), bin the particles into each minimum depending on the shortest distance to the center.
4. This procedure assigns particles to a label from 1 to 12.

**Figure 4:**
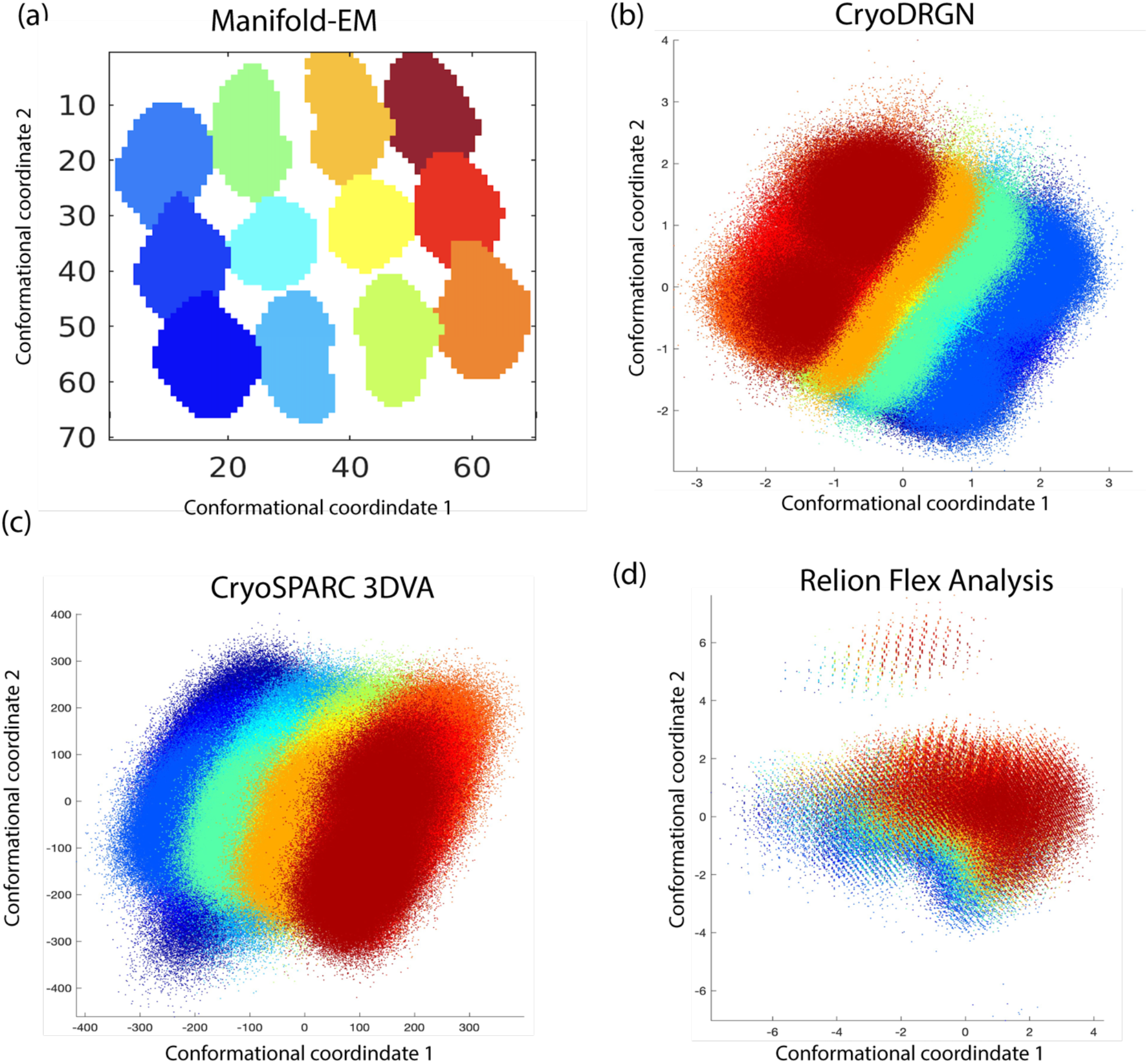
Output conformational landscape colored according to ground truth location (Figure 1(c)) for each of the benchmarked algorithms.

The Accuracy [11] of assigning each minimum ‘n’ is given by:

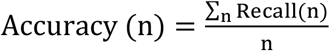

Where TN “True Negatives” is a label where the assignment to the minima ‘n’ is incorrect, TP “True Positive” is a label in the output correctly assigned, and T(n) is the total number of particles in a well-defined region around the minima.

Recall ranges from 0 to 1 (as evident from the definition), where a value of 0 implies no snapshot is correctly assigned to that energy minima and a value of 1 when all snapshots belonging to the energy minima are correctly allocated. The Accuracy of each algorithm is the average recall metric for assignment into all 12 minima.

## Notes

### Competing Interest Statement

The authors have declared no competing interest.

