## Supplementary Information for "Energy Landscapes from Cryo-EM Snapshots: A Benchmarking Study"

#### SUPPLEMENTARY INFORMATION

Raison Dsouza, Ghoncheh Mashayekhi, Roshanak Etmaadpour, Peter Schwander, Abbas Ourmazd*

University of Wisconsin Milwaukee, 3135 N. Maryland Ave, Milwaukee WI 53211, USA

**Supplementary Figures 1-3**

#### Input data and preprocessing steps

We describe the generation of the synthetic cryo-EM snapshots and the steps involved in single-particle cryo-EM image preprocessing. For this purpose, a ribosome-like model with two degrees of freedom entailing rotations of the small subunit about two different axes was simulated (Supplementary figure 1(a)). The 3D density map of an 80S Ribosome at 7A [1] was used as the source for data generation. The ground truth conformations lie in a two-dimensional flat manifold (Supplementary figure 1(b)), which corresponds to the rotation of the lower subunit with respect to the large subunit. We use 180 projection directions along a Great Circle each containing 18000 single-particle images projected along the x-axis. The images are not modified by a microscope contrast transfer function A sample image with and without noise is shown in Supplementary Figures 2(a) and 2(b). The Pixel noise was incorporated via a Gaussian model and particle images are individually normalized to have a unit signal variance on average.

Synthetic cryo-EM snapshots of different conformations were generated with probabilities reflecting a conformational energy landscape consisting of a rectangular 3x4 array of 12 energy regions of different depths (Supplementary figure 1(b)). The 3x4 array was selected to indicate the different extents in conformational change i.e., 6 degrees along conformational coordinate 1 and 4 degrees along conformational coordinate 2. The ground truth energy landscape features modulations of the conformational occupancy to mimic the biomolecule traversing through stable states and transition states. In this regard, we formulated an occupancy distribution via a potential with pockets to imply energy minima. Conformational coordinate 1 was assigned 4 pockets (energy minima) for a corresponding conformational coordinate 2 with three pockets.

#### Conformational landscapes

The data-analytical tools [2-5] utilize powerful machine learning algorithms to determine the number of degrees of freedom exercised by the biomolecule. To assess each algorithm's ability to capture continuous conformational changes, we examined how the input data are distributed of the two leading eigenvectors calculated in every algorithm. Binning the data along these eigenvectors allows us to easily visualize the distribution as histograms. The histograms of the individual conformational coordinates are shown in Supplementary figure 3(a-c). The histograms of conformational coordinates essentially point out the occupancy of each region in the energy landscape. For example, in the ground truth, conformational coordinate 1 shows four highly occupied regions (in energy landscape terms, four minima). Similarly, conformational coordinate 2 shows the three conformational maxima, or equivalently, three energy minima.

To determine the relationship between the different eigenvectors obtained from each method, we plot the snapshots as points on a two-dimensional diagram. Such diagrams called 'Scatter diagrams' enable us to correlate data between different degrees of freedom extracted from the data. Figures 3 (a-d) show the scatter plots of the distribution along each conformational coordinate with the corresponding histograms for each component (along the margin). The histograms of conformational coordinates obtained from Manifold-EM (Supplementary figure 3(a)) show the four energy maxima along with coordinate one and three maxima along with coordinate two. However, the heights of the peaks along conformational coordinate two seemed to bin differently than the ground truth. This leads to some particles being assigned to incorrect energy minima which leads to increased false negatives in the calculation of the accuracy score (See Methods). In cryoDRGN (Supplementary figure 3(b)), the particle distribution is centered along with conformational coordinates one and two, which leads to incorrect assignment of particles into different energy minima regions. This results in an increased standard deviation in the accuracy score between the twelve energy minima. In Relion (Supplementary figure 3(c)), the marginal histograms for the first two principal components are bimodal and the subtle features of the ground truth distribution are lost. This causes a drastic reduction in the accuracy score due to most of the particles being assigned incorrectly. For CryoSPARC 3DVA (Supplementary figure 3(d)), the histograms show an almost featureless histogram for conformational coordinate 1 and monomodal (bell-shaped) distribution for conformational coordinate two. The flat distribution essentially misses out on the subtle features of the ground truth energy landscape.

#### Computational resources

All computations were performed on a CPU cluster with the following specifications: 16 CPU nodes, each consisting of two Deca-core E5-2660 V3 “Haswell”/2.6GHz, and 128GB of memory. Initial orientation recovery was performed using RELION 3.1 for inputs to multibody refinement.

#### Data availability

The ground truth density maps and the synthetic images will be made available from the authors upon reasonable request.

#### Supplementary Tables

| Dataset | Number of Particles | Image size (pixels) | Pixel size (A) | Latent space dimension |
|---|---|---|---|---|
| Simulated | 3240000 | 124 | 2.4 | 2 |

**Supplementary Table 1. The dataset statistics.**

| Relion Multibody | Hits | Recall (%) |
|---|---|---|
| Region 1 | 111411 | 38.55 |
| Region 2 | 11814 | 4.299 |
| Region 3 | 87906 | 30.48 |
| Region 4 | 27727 | 10.61 |
| Region 5 | 8949 | 3.61 |
| Region 6 | 40624 | 15.48 |
| Region 7 | 30824 | 11.72 |
| Region 8 | 7846 | 3.14 |
| Region 9 | 31640 | 12.04 |
| Region 10 | 69794 | 23.93 |
| Region 11 | 26985 | 9.79 |
| Region 12 | 136530 | 46.94 |

**Supplementary Table 2. Recall scores for each energy minimum region obtain from Relion multibody algorithm.**

| cryoSPARC 3DVA | Hits | Recall (%) |
|---|---|---|
| Region 1 | 163349 | 70.65 |
| Region 2 | 75373 | 34.21 |
| Region 3 | 149773 | 64.87 |
| Region 4 | 122002 | 58.21 |
| Region 5 | 51724 | 26.03 |
| Region 6 | 127377 | 60.53 |
| Region 7 | 115827 | 55.05 |
| Region 8 | 47827 | 23.96 |
| Region 9 | 135398 | 64.37 |
| Region 10 | 142466 | 61.04 |
| Region 11 | 53648 | 24.33 |
| Region 12 | 168327 | 72.29 |

**Supplementary Table 3. Recall scores for each energy minimum region obtain from the Cryosparc 3DVA algorithm.**

| cryoDRGN VAE | Hits | Recall (%) |
|---|---|---|
| Region 1 | 188659 | 65.27 |
| Region 2 | 134148 | 48.72 |
| Region 3 | 198849 | 68.94 |
| Region 4 | 176817 | 67.66 |
| Region 5 | 121080 | 48.86 |
| Region 6 | 174829 | 66.65 |
| Region 7 | 180603 | 68.67 |
| Region 8 | 121864 | 48.85 |
| Region 9 | 178956 | 68.10 |
| Region 10 | 200691 | 68.82 |
| Region 11 | 128649 | 46.70 |
| Region 12 | 195169 | 67.11 |

**Supplementary Table 4. Recall scores for each energy minimum region obtain from the Cryodrgn VAE algorithm.**

| **Manifold EM** | **Hits** | **Recall (%)** |
|---|---|---|
| Region 1 | 40189 | 83.11 |
| Region 2 | 34852 | 74.99 |
| Region 3 | 36417 | 78.21 |
| Region 4 | 41721 | 85.24 |
| Region 5 | 36038 | 74.49 |
| Region 6 | 32222 | 71.16 |
| Region 7 | 32873 | 71.62 |
| Region 8 | 35257 | 73.47 |
| Region 9 | 42032 | 83.96 |
| Region 10 | 36714 | 78.24 |
| Region 11 | 36001 | 76.08 |
| Region 12 | 40619 | 80.67 |

**Supplementary Table 5. Recall scores for each energy minimum region obtain from the ManifoldEM algorithm.**

### Supplementary Figures

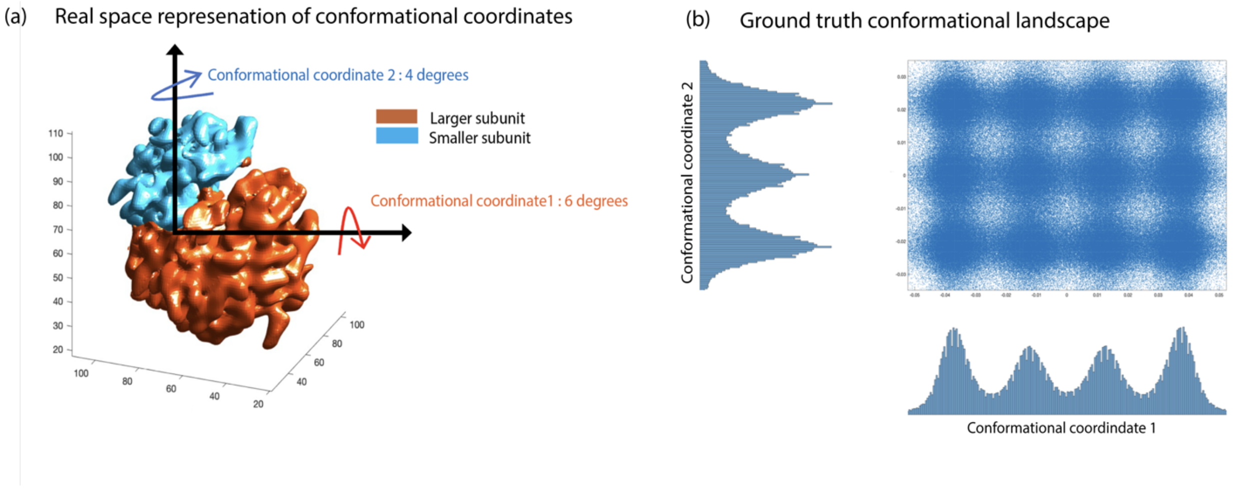

**Supplementary Fig. 1. (a). Sketch of the synthetic model, rotation of the small subunit about the two axes labeled conformational coordinate 1 and 2 respectively. The larger subunit (orange) is fixed. (b). Ground truth conformational landscape for the synthetic model with 12 high occupancy regions, indicated as maxima on the histograms.**

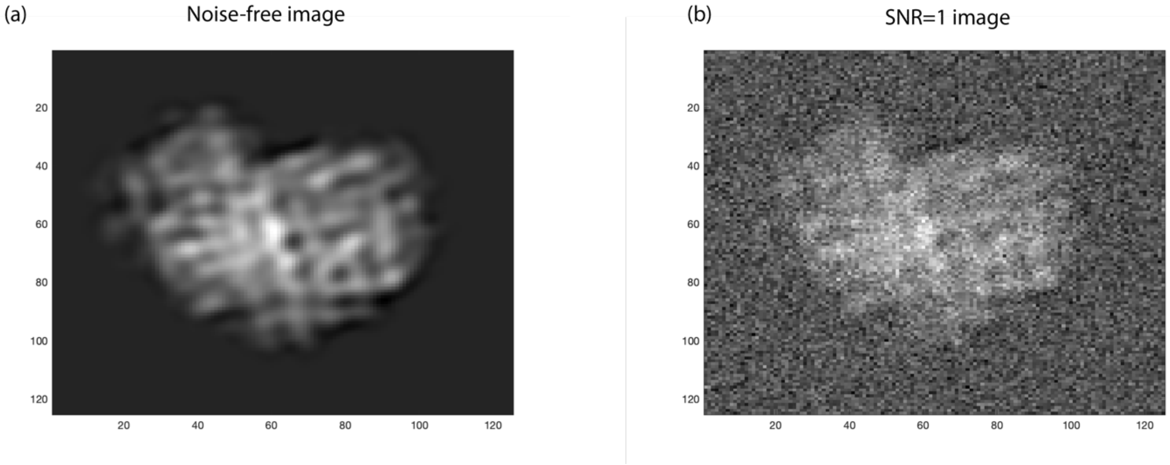

**Supplementary Fig. 2. Images of the synthetic data (a). without pixel noise (b) with pixel noise (SNR=1)**

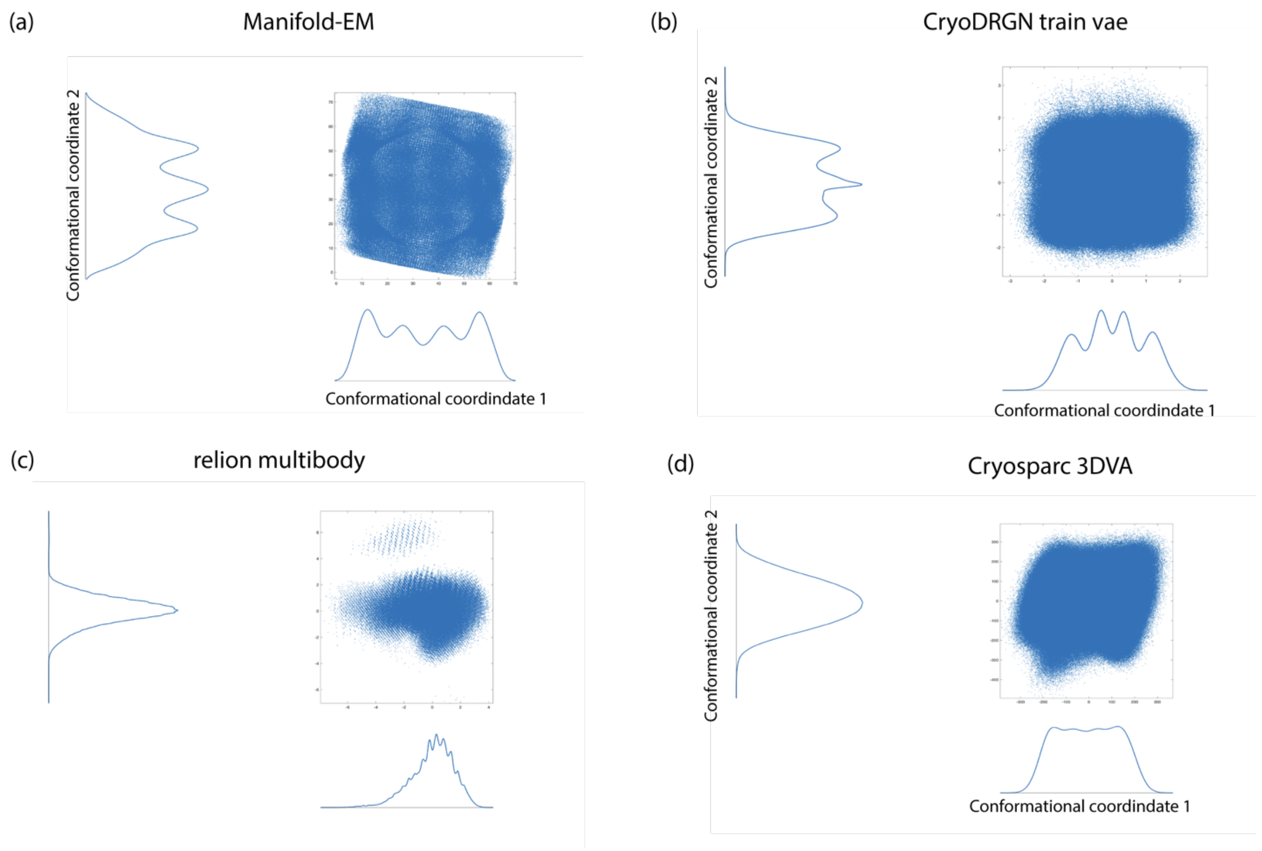

**Supplementary Fig. 3.**

**Scatterplot of the two retrieved conformational coordinates obtained from four algorithms. Also shown are the marginal histograms for each case.**
